# Gene editing preserves visual function in a mouse model of retinal degeneration

**DOI:** 10.1101/624858

**Authors:** Paola Vagni, Laura E. Perlini, Naïg A. L. Chenais, Tommaso Marchetti, Martina Parrini, Andrea Contestabile, Laura Cancedda, Diego Ghezzi

## Abstract

Inherited retinal dystrophies are a large and heterogeneous group of degenerative diseases caused by mutations in various genes. Given the favourable anatomical and immunological characteristics of the eye, gene therapy holds great potential for their treatment. We used a tailored CRISPR/Cas9-based gene editing system to prevent retinal photoreceptor death in the Rd10 mouse model of retinitis pigmentosa. We tested the gene editing tool *in vitro* and then used *in vivo* subretinal electroporation to deliver it to one of the retinas of mouse pups at different stages of photoreceptor differentiation. Three months after gene editing, the treated eye exhibited a higher visual acuity compared to the untreated eye. Moreover, we observed preservation of light-evoked responses both in explanted retinas and in the visual cortex of treated animals. Our study validates a CRISPR/Cas9-based therapy as a valuable new approach for the treatment of retinitis pigmentosa caused by autosomal recessive loss-of-function point mutations.

## Introduction

Retinitis pigmentosa is a group of inherited retinal dystrophies (IRDs) that cause the progressive death of retinal photoreceptors and eventually blindness(Ferrari et al., 2011). The treatment of retinitis pigmentosa is still a major challenge because of the early death of rod photoreceptors and the late onset of the symptoms. Daily vision in humans mainly depends on cone photoreceptors, which in retinitis pigmentosa degenerate only at a late stage: likely because cones metabolically depend on rods, which provide them nutrients(Narayan et al., 2016). Therefore, acting on the principal cause of degeneration, namely at the level of rod photoreceptors, would be an effective therapeutic approach to preserve vision in retinitis pigmentosa. Notably, rod-rich photoreceptor transplantations can halt cone loss in degenerating retinas(Mohand-Said et al., 2000).

Mutations in the β-domain of the phosphodiesterase 6 (*PDE6B*) gene, which hydrolyses cyclic guanosine monophosphate (cGMP) and initiates phototransduction, are among the most commonly identified causes of autosomal recessive retinitis pigmentosa(DANCIGER et al., 1995; McLaughlin et al., 1995). Missense mutations in *PDE6B* lead to photoreceptor death, triggered by the toxic accumulation of cGMP(Ulshafer et al., 1980), and result in a progressive loss of visual function, starting from the peripheral retina and progressing towards the centre. The discovery of naturally occurring mouse models carrying mutations on the *Pde6b* gene(Chang et al., 2002; 2007) has provided a better understanding of the mechanisms underlying retinal degeneration and has prompted the development of new therapies. The retinal degeneration 10 (Rd10) mouse carries an autosomal recessive loss-of-function missense point mutation in the *Pde6b* gene (exon 13; C1678T → R560C), leading to the progressive degeneration of photoreceptor cells. Rd10 mice are particularly useful as an animal model for autosomal recessive retinitis pigmentosa since the slow degeneration of photoreceptor cells recapitulates the time course of the disease in patients.

The first genetic approaches to vision restoration in the Rd10 mouse was based on virus-mediated supplementation of the *Pde6b* gene(Bennett et al., 1996; Jomary et al., 1997; Pang et al., 2008; 2012). Similarly, viral gene transfer therapies have led to promising results for Leber Congenital Amaurosis 2 and some other retinal diseases, as demonstrated by the several ongoing clinical trials(Auricchio et al., 2017). Recently, gene editing tools based on Clustered Regularly Interspaced Short Palindromic Repeats (CRISPRs)-Associated (Cas) Genes have completely revolutionised gene therapy(Heidenreich and Zhang, 2016). The Cas9 nuclease utilises a guide RNA (gRNA) to induce DNA Double-Strand Breaks (DSBs) at a precise location in the target genomic site. CRISPR/Cas9 system is easily tuneable, versatile, and enables the precise correction of genetic defects directly on the patient genome. The CRISPR/Cas9 system can either be used to disrupt the target gene by Non-Homologous End-Joining (NHEJ) of DSBs or to edit the target gene by Homology Directed Repair (HDR) in the presence of a DNA donor sequence (repair template). Importantly, the expression of the CRISPR/Cas9 system is only needed for the relatively short period necessary to correct the genetic mutation (a few days, rather than continuously as in the case of gene supplementation therapies)(Ran et al., 2013).

In this study, we designed a CRISPR/Cas9 gene editing system that can repair the genetic mutation in the Rd10 mouse model taking advantage of the increased activity of the HDR mechanism in dividing progenitor cells(Saleh-Gohari and Helleday, 2004). We tested the efficiency of the designed approach first *in vitro* and then *in vivo*. To demonstrate the phenotype reversal, we performed behavioural and electrophysiological analysis on treated and control mice. Overall, the treated mice retained 50 % of the normal visual acuity even three months after the treatment.

## Results

### gRNA screening and high editing efficiency of the CRISPR-Cas9 vector in cell culture

The efficiency of different gRNAs in inducing HDR-mediated editing of a specific genomic locus can be very different, ranging from 0.7 to 30 %(Cong et al., 2013; Ran et al., 2013). Therefore, as a first step in the development of the gene editing system, we designed and screened different gRNAs for their ability to induce CRISPR/Cas9-mediated editing of the *Pde6b* gene. We selected three candidate gRNAs and screened them in mouse Neuro 2A (N2A) cells to determine which one was the best at targeting the sequence coding for WT *Pde6b*. We transfected N2A cells with a single plasmid, containing Cas9, one of the three gRNAs, and the green fluorescent protein (GFP), along with a DNA single-stranded oligonucleotide (ssODN) repair template specific for each gRNA, containing flanking sequences of 100 bp on each side of the insertion site that were homologous to the target region. The gRNA #1 and gRNA #3 mapped upstream and downstream to the Rd10 locus, while the gRNA #2 mapped directly on it (**Fig. 1a**). Each repair template for HDR-mediated editing was designed to edit the genomic DNA (gDNA) sequence at the Rd10 locus and simultaneously remove an adjacent cutting site for the restriction enzyme BanI (by introducing a silent mutation), allowing the assessment of the editing efficiency by BanI restriction analysis (**Fig 1a)**. Moreover, each repair template also carried a second specific silent mutation in the PAM sequence of the corresponding gRNA to avoid further Cas9-mediated cutting on the edited genomic sequence (**Fig. 1a**). One day after transfection, we isolated GFP-expressing cells by fluorescent activated cell sorting (FACS), extracted the gDNA and PCR-amplified a 700 bp fragment containing the *Pde6b* target region with primers mapping outside the ssODN homology arm sequence (**Fig. 1a**). After BanI digestion and agarose-gel electrophoresis, the edited DNA appeared as a single uncut band (700 bp), while non-edited DNA was digested in two fragments (230 and 470 bp). The quantification of the percentage of edited versus non-edited DNA for each gRNA showed that gRNA #2 had the highest editing efficiency and was the best performing gRNA (**Fig. 1b,c**). Based on this result, we next designed for the final editing tool gRNA #4 which differ from gRNA #2 only in a single base pair (**Fig. 1d**), corresponding to the C to T mutation found in the mutated *Pde6b* gene of Rd10 mice. We further verified gRNA #4 editing efficiency in neural stem cells (NSC) derived from Rd10 homozygous pups. The transfected Rd10 cells were selected for GFP-expression with FACS and the editing efficiency was evaluated by BanI restriction assay, as described above for N2A cells. We found a mean (± s.d., *n* = 3) net editing efficiency of 39.3 ± 6.4 % in NSC harbouring the Rd10 mutation (**Fig. 1e,f**). These data indicate that the selected gRNA #4 efficiently targets the Rd10 mutation in the *Pde6b* gene and that the correct sequence can be restored with high efficiency by CRISPR/Cas9-mediated HDR editing.

**Figure 1.**
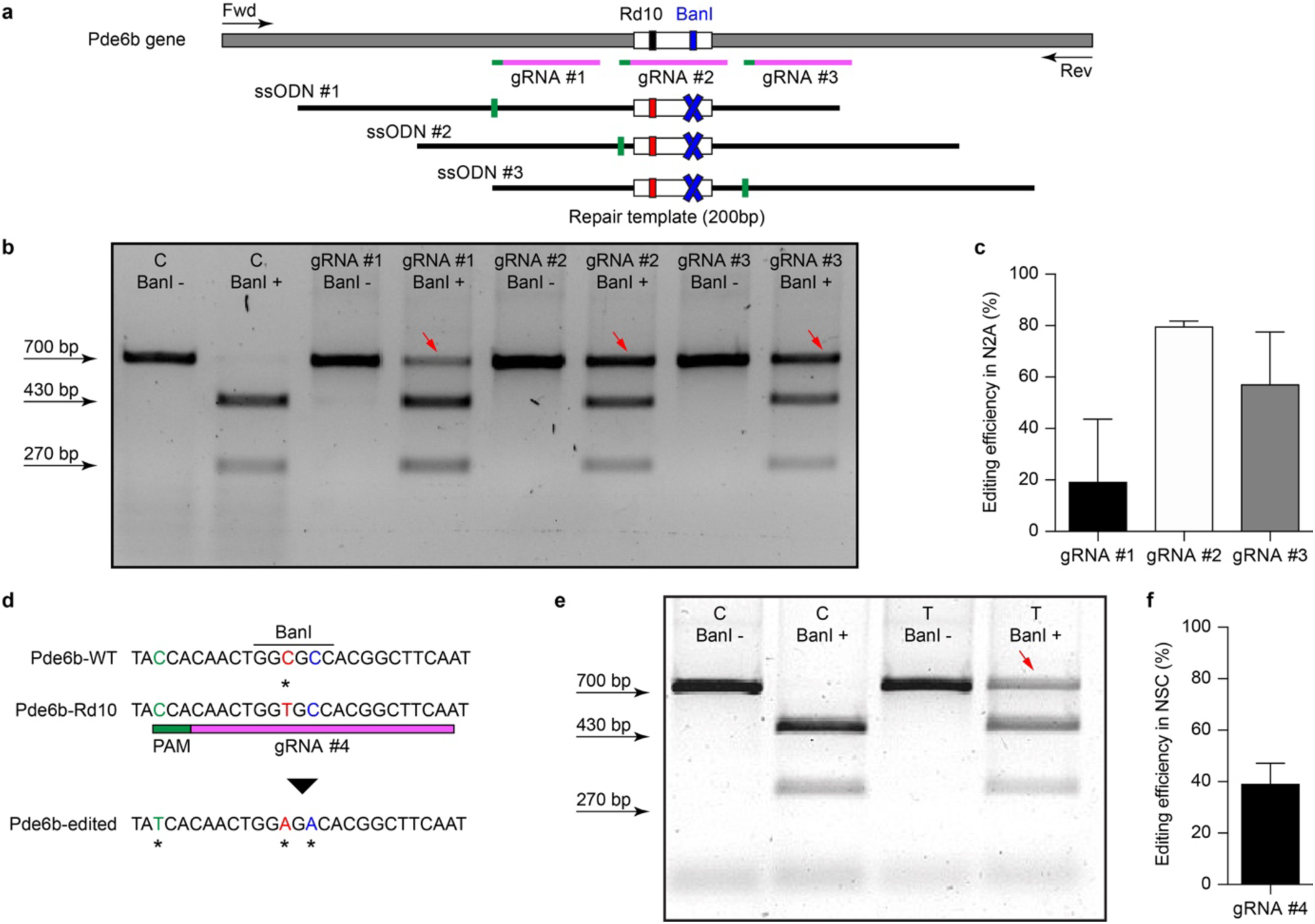
Screening of gRNAs targeting the Rd10 locus. **a**, Schematic representation (not in scale) of the mouse *Pde6b* gene showing the position of the three gRNAs tested (in magenta with green PAM sequence), the ssODN repair templates (black), and the PCR primers used for screening (arrows). The white rectangle represents the target editing region with Rd10 mutation (black/red) and the BanI cutting site (blue). Each ssODN also carries a silent mutation in the corresponding gRNA PAM sequence (green). **b,** Representative example of an agarose gel electrophoresis of the BanI restriction assay from transfected (T) and control (C) mouse N2A cells. Unedited DNA is cut in two fragments by BanI digestion (470 and 230 bp), while edited DNA is not cut by the restriction enzyme (700 bp band, red arrows). **c**, Quantification of the mean (± s.d., *n* = 2) editing efficiency for the three gRNA in N2A cells. **d,** Schematic representation of editing strategy for gRNA #4 targeting the Rd10 mutation. The HDR strategy was designed to edit the DNA sequence (in red), while introducing a silent mutation in the cutting sequence for BanI (in blue). A second silent mutation in the PAM sequence of the gRNA (in green) is included in the repair template in order to avoid further Cas9-mediated cutting on the edited genomic sequence. **e,** Representative example of an agarose gel electrophoresis of the BanI restriction assay for gRNA #4 transfected (T) and control (C) NSC from Rd10 mice. The red arrow indicates the edited DNA that is resistant to BanI digestion. **c**, Quantification of the mean (± s.d., *n* = 3) editing efficiency for gRNA #4 in Rd10 NSC.

### Efficient delivery of a reporter gene to photoreceptor cells by in vivo electroporation

The efficiency of electrotransfer depends on various factors such as the cell size, the parameters of the electric pulses, and the phase of the cell cycle. The latter has to be taken into account especially when interested in targeting the highest number of cells and in exploiting the HDR mechanism to achieve gene editing. In order to obtain the highest number of transfected cells without inducing eye defects, we performed pilot experiments to optimise the electroporation protocol and select the best timing for delivery. Although electroporation immediately after birth is potentially more efficient, it can result in eye damage: in our hand, postnatal day (P) 1 electroporation resulted in more than 50 % of the pups bearing eye defects as adults, while this percentage was reduced at 40 % by performing electroporation at P3 (at the peak of the photoreceptor proliferation curve). For this reason, P3 was selected for the *in vivo* experiments (**Fig. 2a,** dashed line). Moreover, several groups reported efficient retinal electroporation in neonatal mice using five pulses of 50 ms at 80 V (1 Hz)(Matsuda and Cepko, 2004; de Melo and Blackshaw, 2011), but we found this protocol to cause eye defects (possibly affecting visual functions) in 40 % of the adult mice when the electroporation was performed at P3. Thus, based on previous observations in cell cultures(Bureau et al., 2000), we tested a different protocol (**Fig. 2b**) consisting of two short poring pulses at high voltage (5 ms, 100 V, 0.1 Hz) followed by five long transfer pulses at a lower voltage (50 ms, 30 V, 1 Hz). Applying this improved protocol, we obtained an electroporation efficiency comparable to the standard protocol, whereas the number of pups bearing eye defects when adults decreased from 40 to 5 %; those animals were excluded from the experiments. Also, the use of a conductive gel between the electrode plate and the tissue increased the conductivity and avoided burning marks on the cornea. In order to assess the efficiency of electroporation in targeting the photoreceptor cell progenitors *in vivo*, we delivered a plasmid coding for eGFP to the subretinal space of Rd10 mouse pups with two consecutive subretinal injections followed by electroporation at P3 (**Fig. 2c**). The image sequence shows that eGFP was expressed at all the different stages of retinal development at which the retinas were isolated: P5, P10, and P15 (*n* = 6 at each time point). At P5, most of the expressing cells were confined to the ventricular zone (VZ), where the photoreceptor progenitors proliferate. At P10, the cells started to migrate towards the photoreceptor layer, which they finally reached by P15. The electroporation targets mostly photoreceptors due to their proximity to the injection site, but it is not completely specific to this cell type; indeed, we observed some bipolar and ganglion cells expressing eGFP. This eventuality does not represent a concern for the outcome of the therapy since the targeted gene is expressed specifically in rod photoreceptors. To analyse the extension of the electroporated zone, we prepared wholemount retinas from the treated mice (**Fig. 2d**). In a few cases (2 out of 6) the two consecutive injections per eye resulted in two electroporated areas and all the other cases in one area only, with a single area covering up to 25 % of the retina. The localisation of the electroporated cells depends on the orientation of the electric field to the injection site at the moment of the electroporation, which is challenging to control in an animal as small as the mouse, especially at this young age.

**Figure 2.**
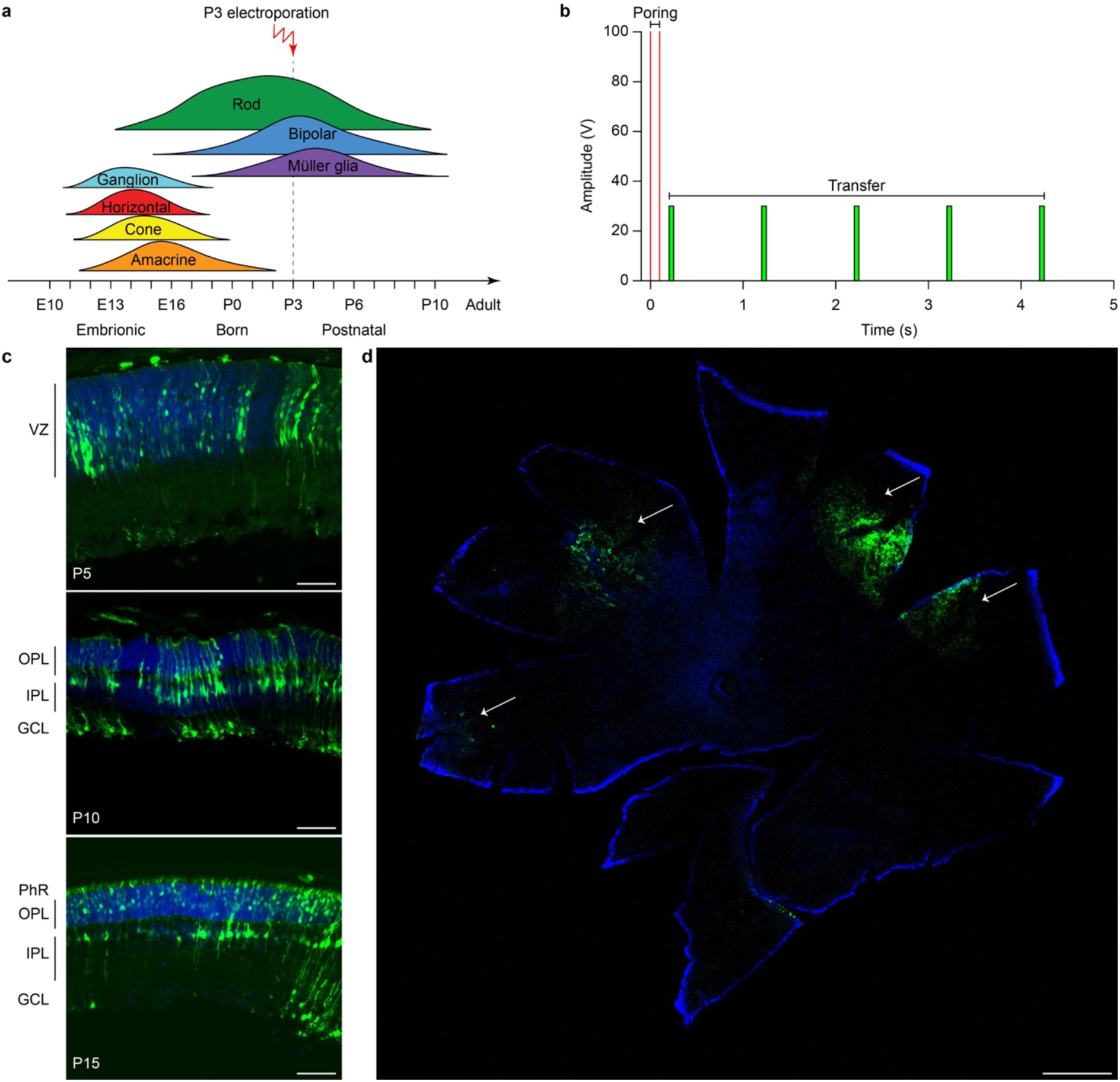
Electroporation of photoreceptor progenitor cells *in vivo* at P3. **a**, The graph shows the proliferation period for all the retinal cell types Sketch redrawn from(Zhang et al., 2011). The proliferation of rod photoreceptors has a peak at birth (P0-P3) and continues until P10. The electroporation was performed at P3. **b,** Schematic representation of the protocol used for electroporation. Two high voltage poring pulses (5 ms, 100 V, 0.1 Hz) are followed by five low voltage transfer pulses (50 ms, 30 V, 1 Hz). **c,** Retinal sections from Rd10 mice electroporated at P3 and collected at different time points, starting from the top: P5, P10, and P15. The scale bar is 60 µm. On the side, the ventricular zone (VZ), the photoreceptor layer (PhR), the outer plexiform layer (OPL), the inner plexiform layer (IPL), and the ganglion cell layer (GCL) are reported. **d,** Representative wholemount retina electroporated at P3 and collected at P10 illustrating the spread of the electroporation (white arrows). The scale bar is 500 µm.

### Significant editing efficiency in vivo of the CRISPR-Cas9 editing tool

Evaluating editing efficiency in whole retinas *in vivo* is a more challenging task than *in vitro* due to the presence of a mixed population of transfected and non-transfected cells. To this aim, we developed a sensitive droplet digital PCR (ddPCR) assay with two fluorescent probes specific for the edited and unedited alleles (**Fig. 3a**). Rd10 pups were electroporated at P3 with plasmids encoding eGFP, Cas9, gRNA #4, together with the ssODN repair template. Sham electroporation of control retinas in Rd10 pups at P3 was performed by omitting the gRNA. Three days after electroporation, we extracted the gDNA from whole retinas and analysed the editing efficiency at the *Pde6b* gene by ddPCR. We found that the mean (± s.d., *n* = 10) *in vivo* editing efficiency in Rd10 treated retinas was 0.22 ± 0.14 % and significantly different from Rd10 sham retinas (p < 0.05, unpaired t-test) that however showed a low but detectable background (0.057 ± 0.050 %, *n* = 6) in the assay (**Fig. 3b)**. Although *in vivo* editing efficiency appeared much lower than *in vitro*, this represents an underestimation because the assay was conducted on gDNA extracted from whole retinas that contained only a relatively small percentage of transfected cells. Moreover, treated retinas showed variable degrease of editing, likely due to a difference in electroporation efficiency. However, even a few functional photoreceptors can make a large difference when it comes to visual performance(MacLaren et al., 2006; Tucker et al., 2011).

**Figure 3.**
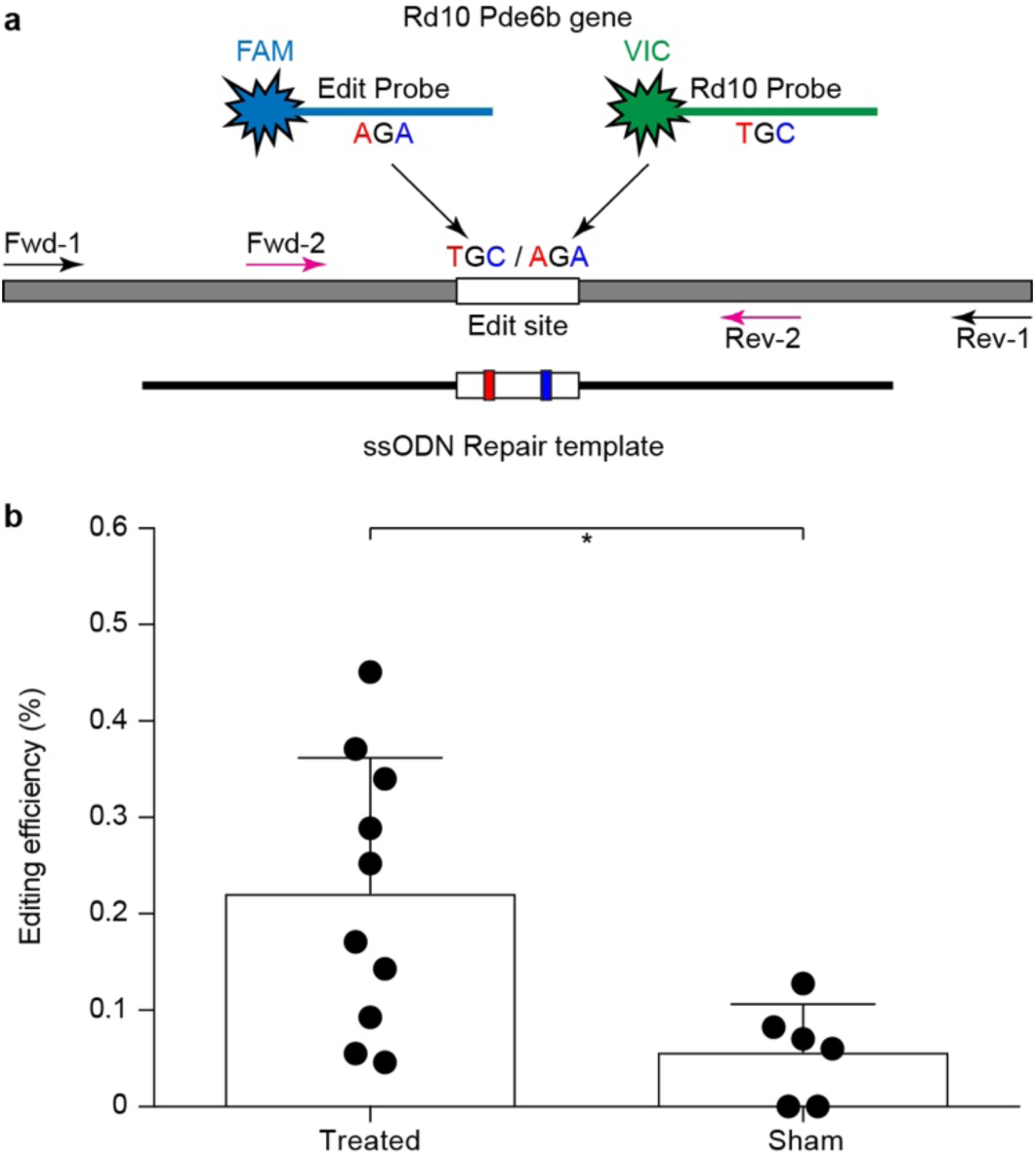
Editing efficiency in vivo. **a**, Schematic representation of the ddPCR assay used to quantify *in vivo* editing efficiency. The *Pde6b* gene is in grey and the ssODN repair template in black (not in scale). The white rectangle represents the target editing region. Black and magenta arrows indicate primers pairs used for the nested-ddPCR. Red and blue letters indicate base mismatches detected by the two specific fluorescent probes for the edited and unedited alleles (blue and green respectively). **b**, Quantification of the mean (± s.d.) percentage of editing in Rd10 treated retinas.

### Gene editing preserves microelectroretinograms ex vivo at P60

To verify whether the extent of gene editing can translate to improved retinal functionality, we recorded the microelectroretinograms (µERGs) from explanted retinas of P60 Rd10 mice that were electroporated at P3 (**Fig. 4a**). Untreated Rd10 mice were used as control. Previous results show that Rd10 retinas are completely degenerated(Jae et al., 2013) and stop consistently responding to light stimulation at P60(Stasheff et al., 2011). We recorded simultaneously from all the electrodes of a multielectrode array (MEA) while stimulating using green light pulses (4 ms, 0.5 mW mm^-2^). In **Fig. 4b,** we present a representative µERG response from a treated retina, as the average over ten sequential stimulations delivered at 1 Hz of repetition rate. The a-wave peak amplitudes in Rd10 treated retinas are significantly higher (p < 0.0001, unpaired t-test) than Rd10 untreated retinas (**Fig. 4c**). This result supports our hypothesis that the functionality of the retina is preserved in Rd10 treated mice.

**Figure 4.**
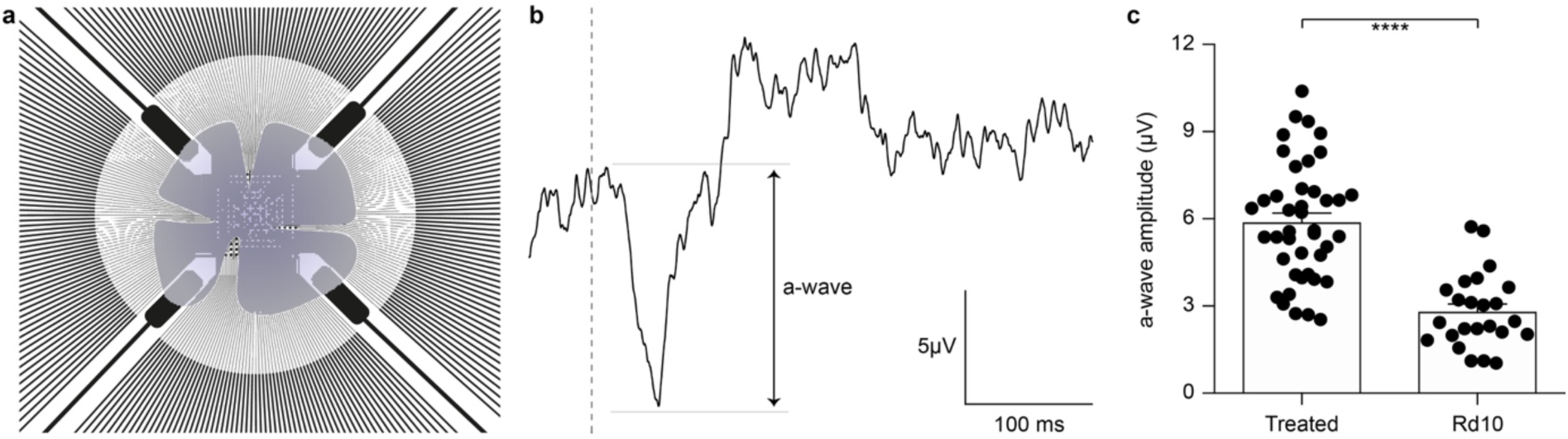
Preservation of microelectroretinograms at P60. **a**, Schematic representation of the experiment. The retina was dissected and placed on a transparent MEA with the retinal ganglion cell side down in contact with the electrodes. The retina was stimulated using green light pulses coming from the bottom. **b,** Representative recording from a Rd10 retina treated at P3. **c,** Quantification of the mean (± s.e.m.) amplitude of the a-wave in the two experimental groups: Rd10 treated (5.89 ± 0.31, *n* = 42 channels from 2 retinas) and Rd10 untreated (2.81 ± 0.26, *n* = 27 channels from 2 retinas) retinas.

### Gene editing preserves visual acuity in vivo until P90

Next, we verified whether our results *ex vivo* could effectively lead to functional improvement of vision *in vivo*. To this end, we assessed the visual acuity of Rd10 unilaterally treated (at P3), Rd10 unilaterally sham-treated (at P3), Rd10 untreated, and WT adult mice using a behavioural assay. The optomotor test, which measures the integrity of the subcortical visual pathways, uses the amplitude of the optomotor reflex to evaluate the visual acuity of rodents (**Fig. 5a**). In particular, it allows the distinction between right eye-driven and left eye-driven responses, measuring the visual threshold of each eye independently(DOUGLAS et al., 2005). At P30 (about 1 month after treatment), in Rd10 mice, the treated eye (**Fig. 5b**, white circles) showed a higher visual acuity compared to the paired untreated eye (treated 0.24 ± 0.01 C/°, untreated 0.14 ± 0.01 C/°; *n* = 71, p < 0.0001, Mann-Whitney test). Conversely, the Rd10 sham-treated eye (light grey circles) did not show any improvement compared to the paired untreated eye (sham 0.11 ± 0.01 C/°, untreated 0.13 ± 0.01 C/°; *n* = 20, p = 0.2306, unpaired t-test). In both WT (left 0.41 ± 0.01 C/°, right 0.39 ± 0.01 C/°; *n* = 32, p = 0.1009, Mann-Whitney test; black dots) and Rd10 (left 0.12 ± 0.01 C/°, right 0.11 ± 0.01 C/°; *n* = 47, p = 0.4908, unpaired t-test; dark grey dots) mice no difference was detected between the left and right eyes (**Fig. 5b**). Measures of the optomotor reflex (**Fig. 5c**) demonstrated that the average visual acuity in the treated eyes of Rd10 mice is significantly higher than the average visual acuity of both Rd10 mice (p < 0.0001; One Way ANOVA, Tukey’s multiple comparisons test) and sham-treated eyes in Rd10 mice (p < 0.0001; One Way ANOVA, Tukey’s multiple comparisons test). Sham-treated eyes have a visual acuity not statistically different from Rd10 mice’s eyes (p = 0.9849; One Way ANOVA, Tukey’s multiple comparisons test). Since it was not measured in dark-adapted conditions, the outcome of the optomotor test is essentially related to the integrity of cone cells and direction-selective retinal ganglion cells. However, the visual acuity measured with this test is reportedly decreasing in Rd10 mice, already starting from P30(Prusky et al., 2004), which matches our data from control and sham mice. We can thus attribute any further preservation of visual acuity to a protective effect of the treatment.

**Figure 5.**
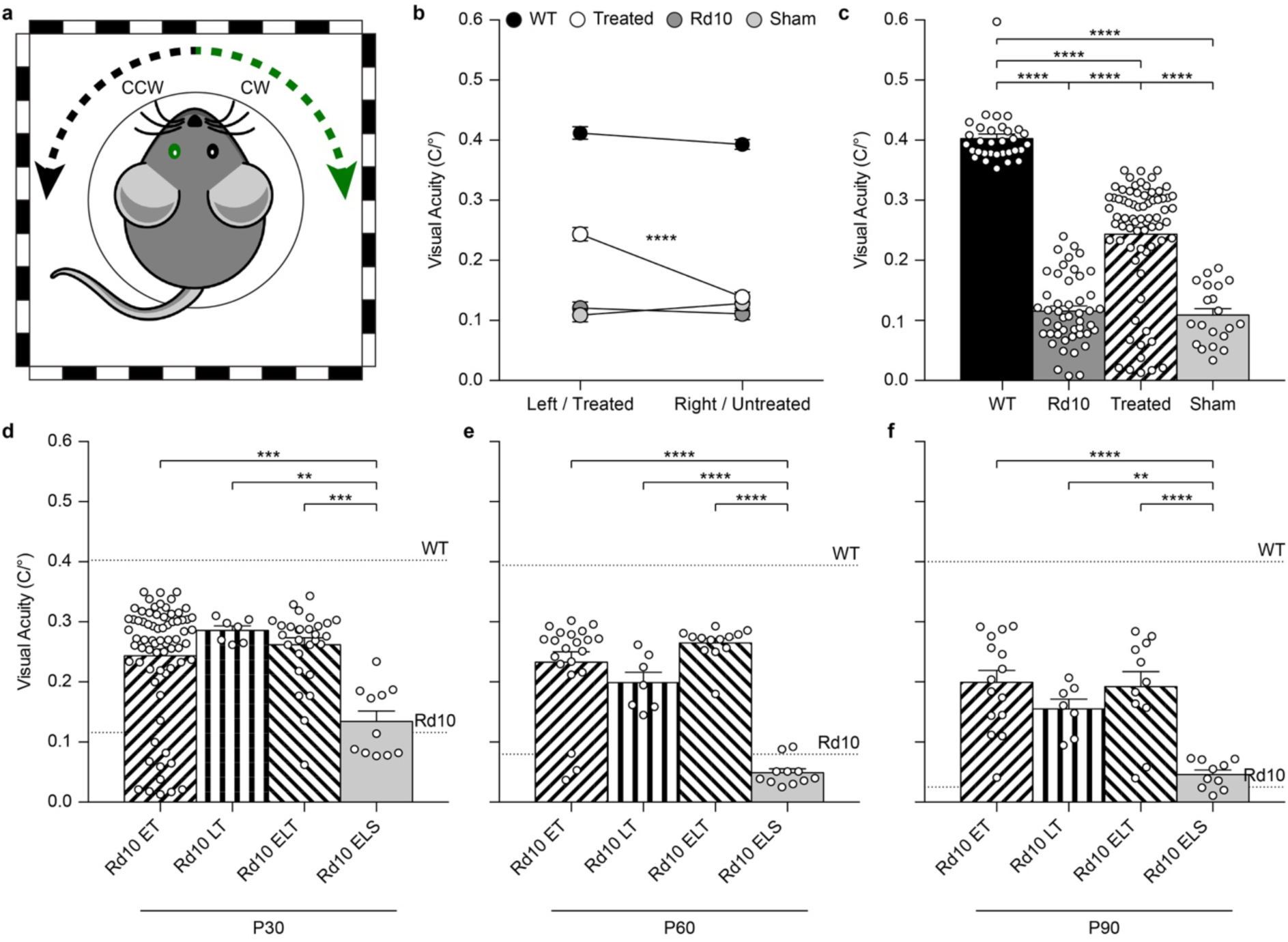
Preservation of the visual acuity in Rd10 treated mice. **a**, For each mouse, both the clock-wise (CW, left eye) and counter clock-wise (CCW, right eye) responses were assessed. The sketch represents a mouse with treatment (green) in the left eye (corresponding to the CW response). **b**, Mean (± s.e.m.) visual acuity in WT mice (black circles), untreated Rd10 mice (dark grey circles), Rd10 treated mice (white circles), and Rd10 sham-treated mice (light grey circles). **c**, Statistical comparison among the 4 groups (p < 0.0001, One Way ANOVA): WT (0.40 ± 0.01 C/°, *n* = 32, averaged left and right responses), Rd10 (0.12 ± 0.01 C/°, *n* = 47, averaged left and right responses), Rd10 treated (0.24 ± 0.01 C/°, *n* = 71), and Rd10 sham (0.11 ± 0.01 C/°, *n* = 20). **d**, Statistical comparison (p < 0.001, One Way ANOVA) of the mean (± s.e.m.) visual acuity among Rd10 ET (0.24 ± 0.01 C/°, *n* = 71), Rd10 LT (0.29 ± 0.01 C/°, *n* = 7), Rd10 ELT (0.26 ± 0.01 C/°, *n* = 28), and Rd10 ELS (0.13 ± 0.02 C/°, n =11) at P30. **e**, Statistical comparison (p < 0.0001, One Way ANOVA) of the mean (± s.e.m.) visual acuity among Rd10 ET (0.23 ± 0.02 C/°, *n* = 21), Rd10 LT (0.20 ± 0.02 C/°, *n* = 7), Rd10 ELT (0.26 ± 0.01 C/°, *n* = 13), and Rd10 ELS (0.05 ± 0.02 C/°, *n* = 11) at P60. **f**, Statistical comparison (p < 0.0001, One Way ANOVA) of the mean (± s.e.m.) visual acuity among Rd10 ET (0.20 ± 0.02 C/°, *n* = 15), Rd10 LT (0.15 ± 0.02 C/°, *n* = 7), Rd10 ELT (0.20 ± 0.03 C/°, *n* = 11), and Rd10 ELS (0.05 ± 0.02 C/°, *n* = 10) at P90. In panels **c-f**, each circle represents a single mouse.

We next investigated whether the same treatment could be effective at a later stage of photoreceptor differentiation. To this end, we treated mice at P8 (**Supplementary Fig. 1a**), approximately at the end of the progenitor cells proliferation curve(Zhang et al., 2011). Electroporation at P8 did not result in any eye damage. After the electroporation at P8, the eGFP fluorescence could be detected in retinas of P10 and P15 mice (**Supplementary Fig. 1b,c**).

**Supplementary Figure 1.**
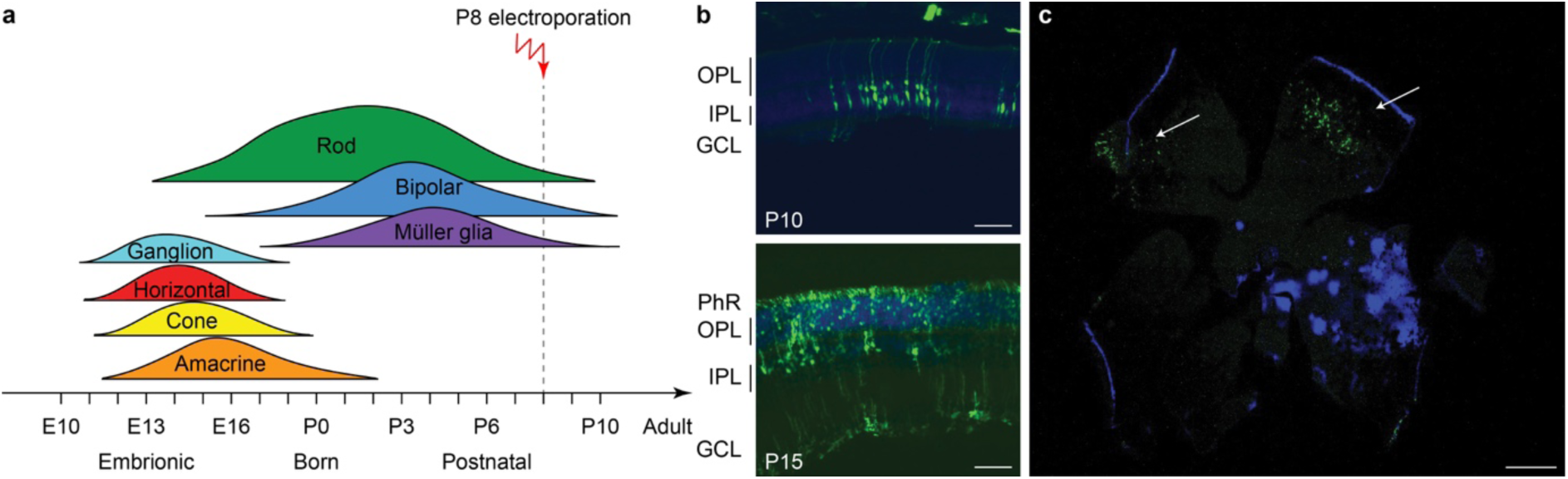
Electroporation *in vivo* of photoreceptor progenitor cells at P8. **a**, Sketch of the late electroporation time point (P8), at the end of the proliferation period. **b,** Retinal sections from mice electroporated at P8. The sections were collected at two time points after electroporation: P10 and P15. The scale bar is 60 µm. On the side, the photoreceptor layer (PhR), the outer plexiform layer (OPL), the inner plexiform layer (IPL), and the ganglion cell layer (GCL) are reported. **c,** Wholemount retina electroporated at P8 and collected at P15, illustrating the spread of the electroporation (white arrows). The scale bar is 500 µm.

Based on this result, we assessed the impact of the period of treatment on the optomotor reflex. We compared the optomotor reflex responses of Rd10 mice upon electroporation at P3 (Rd10 Early Treated, ET), P8 (Rd10 Late Treated, LT), or P3 and P8 (Rd10 Early/Late Treated, ELT). The first treatment corresponds to the peak of the rod’s proliferation curve, the second one to the end of the curve, and the last treatment to the combination of the two (**Fig. 5d**). As for the Rd10 ET mice (**Fig. 5b**), also for the Rd10 LT (**Supplementary Fig. 2a**) and the Rd10 ELT (**Supplementary Fig. 2b**), the visual acuity measured in the treated eyes was significantly higher than the visual acuity of the paired untreated eyes. Conversely, in Rd10 Early/Late Sham (Rd10 ELS) treated mice the visual acuity was not different between injected and not injected eyes (**Supplementary Fig. 2c**).

**Supplementary Figure 2.**
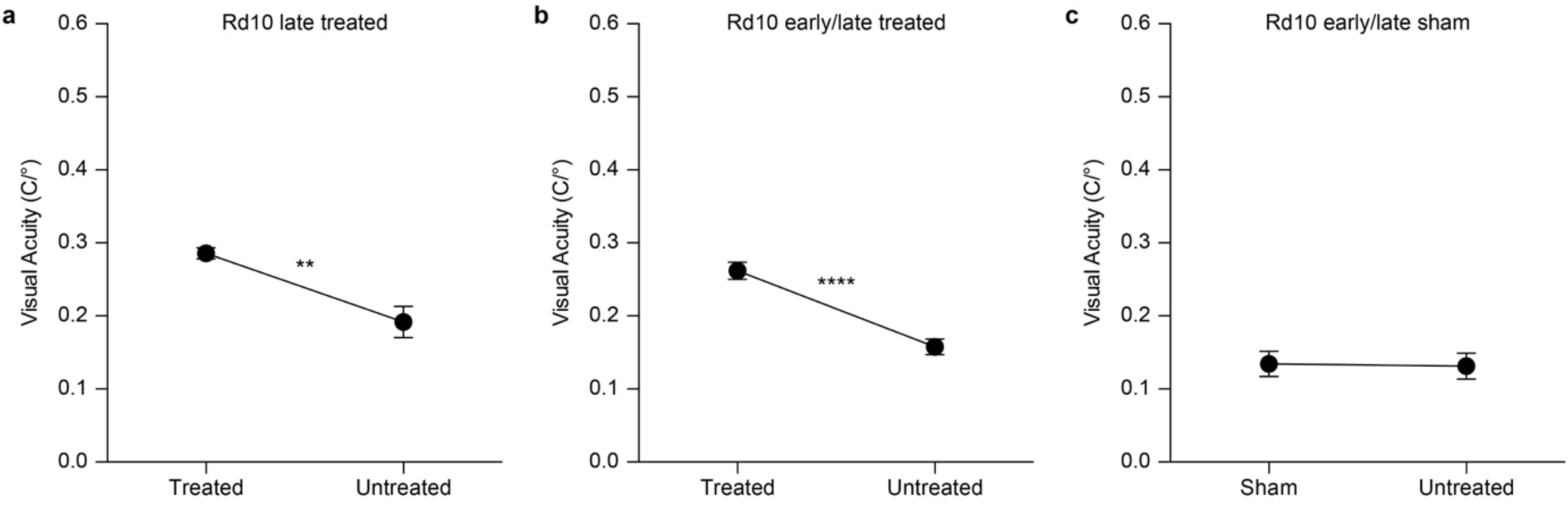
Optomotor reflex upon electroporation at P8. **a**, Mean (± s.e.m.) visual acuity in Rd10 mice treated at P8 (Rd10 LT; treated eye 0.29 ± 0.01 C/°, untreated eye 0.19 ± 0.02 C/°; *n* = 7, p < 0.01, unpaired t-test). **b**, Mean (± s.e.m.) visual acuity in Rd10 mice treated at P3 and P8 (Rd10 ELT; treated eye 0.26 ± 0.01 C/°, untreated eye 0.16 ± 0.01 C/°; *n* = 28, p < 0.0001, unpaired t-test). **c**, Mean (± s.e.m.) visual acuity in Rd10 mice sham treated at P3 and P8 (Rd10 ELS; sham eye 0.13 ± 0.02 C/°, untreated eye 0.13 ± 0.02 C/°; *n* = 11, p = 0.8992, unpaired t-test).

Last, we verified the long-term preservation of visual acuity by repeating the optomotor test at P60 and P90. Interestingly, while the visual acuity dropped drastically in untreated and sham-treated Rd10 mice at P60, it did not show a significant decline in any of the treated groups (**Fig. 5e**). At P90, the visual acuity eventually decreased also in treated mice, but overall all the treated groups retained about 50 % of their initial value (**Fig. 5f**). This result shows a preserved functionality of subcortical visual pathways up to 3 months in treated mice at both P3 and P8.

### Gene editing preserves flash-evoked cortical responses at P90

To assess the functionality of the retino-cortical visual pathway, we recorded visually-evoked cortical potentials (VEPs) from both hemispheres upon flash stimulation. In **Fig. 6a,** we show a representative trace for each experimental group. For treated and sham-treated Rd10 mice, the representative recordings are relative to the cortex contralateral to the injected eye, since in the mouse the majority of the projections decussate at the optic chiasm(Coleman et al., 2009). Since we cannot exclude completely the input coming from the untreated eye (ipsilateral projection), we compared the results of the treated mice with the ones from Rd10 and sham-treated animals to isolate the contribution of the therapy. The mean prominence of the response’s peaks was computed. At P90, Rd10 animals show a complete retinal degeneration with very few spared photoreceptors(Gargini et al., 2007; Pennesi et al., 2012). Accordingly, we observed an almost flat response in untreated (Rd10) and sham-treated (Rd10 ES and Rd10 ELS) mice. Conversely, when recording from all the treated groups (Rd10 ET, Rd10 LT, and Rd10 ELT), we observed preservation of the peak prominence in the visual response of the contralateral cortex (**Fig. 6b**). This is indicative of a preserved functionality of cortical visual pathways (p < 0.0001; Kruskal-Wallis, Dunns multiple comparison test). In the ipsilateral cortex (**Fig. 6c**), the only significant difference was between WT and Rd10 (p < 0.01; Kruskal-Wallis, Dunns multiple comparison test).

**Figure 6.**
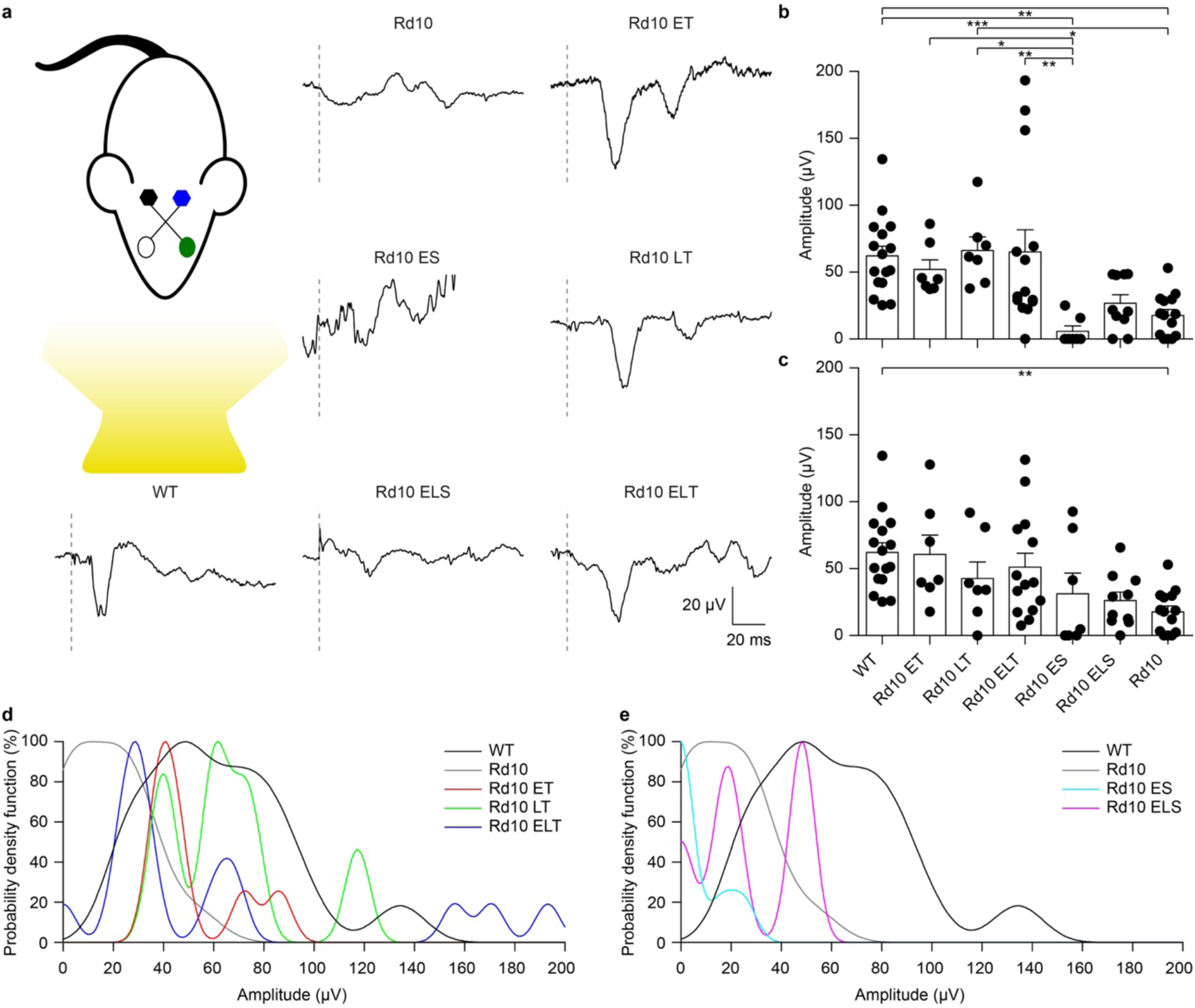
Preservation of visually evoked potentials at P90. **a**, Sketch of the recording setup in which the cortex contralateral to the treated eye (green) is in black, while the ipsilateral is in blue. Representative VEP response for each experimental group. The grey dashed lines are the occurrence of the flash. For treated (Rd10 ET, Rd10 LT, and Rd10 ELT) and sham-treated (Rd10 ES and Rd10 ELS) mice, the traces are from the contralateral cortex, while for WT and Rd10 mice the responses of the two cortices were averaged. **b,** Mean (± s.e.m.) contralateral peak amplitude for all the experimental groups: WT (91.7 ± 11.1 µV, *n* = 16), Rd10 ET (62.4 ± 9.4 µV, *n* = 7), Rd10 LT (73.1 ± 10.8 µV, *n* = 7), Rd10 ELT (71.7 ± 15.2 µV, *n* = 14), Rd10 ES (13.7 ± 8.9 µV, *n* = 7), Rd10 ELS (34.4 ± 7.1 µV, *n* = 10), and Rd10 (23.8 ± 5.5 µV, *n* = 14). **c,** Mean (± s.e.m.) ipsilateral peak amplitude for all the experimental groups: WT (91.7 ± 11.1 µV, *n* = 16), Rd10 ET (69.6 ± 14.4 µV, *n* = 7), Rd10 LT (49.8 ± 12.8 µV, *n* =7), Rd10 ELT (96.6 ± 20.7 µV, *n* = 14), Rd10 ES (48.0 ± 21.9 µV, *n* = 7), Rd10 ELS (30.8 ± 7.0 µV, *n*= 10), and Rd10 (23.8 ± 5.5 µV, *n* = 14). In **b** and **c**, for WT and Rd10 mice the responses of the two cortices were averaged before computing the peak amplitude; therefore, they are equal. **d,** Probability density functions fitted using a Kernel distribution of the contralateral response for the treated groups (Rd10 ET, Rd10 LT, and Rd10 ELT) compared to WT and Rd10 mice. **e,** Probability density functions fitted with a Kernel distribution of the contralateral response for the sham-treated groups (Rd10 ES and Rd10 ELS) mice compared to WT and Rd10 mice.

Finally, we further compared the scaled probability density functions (pdf, **Supplementary Fig. 3**) of the VEP prominences in the treated (**Fig. 6d**) and sham-treated (**Fig. 6e**) mice to the ones of WT and Rd10 mice. In WT mice, the pdf was broadly centred at 50 µV. All of the treated groups had a distribution that appeared to be concentrated around 50 µV and narrower than the one of WT mice. In contrast, sham-treated and control groups distributions are skewed towards 0, highlighting the higher amount of non-responding mice.

**Supplementary Figure 3.**
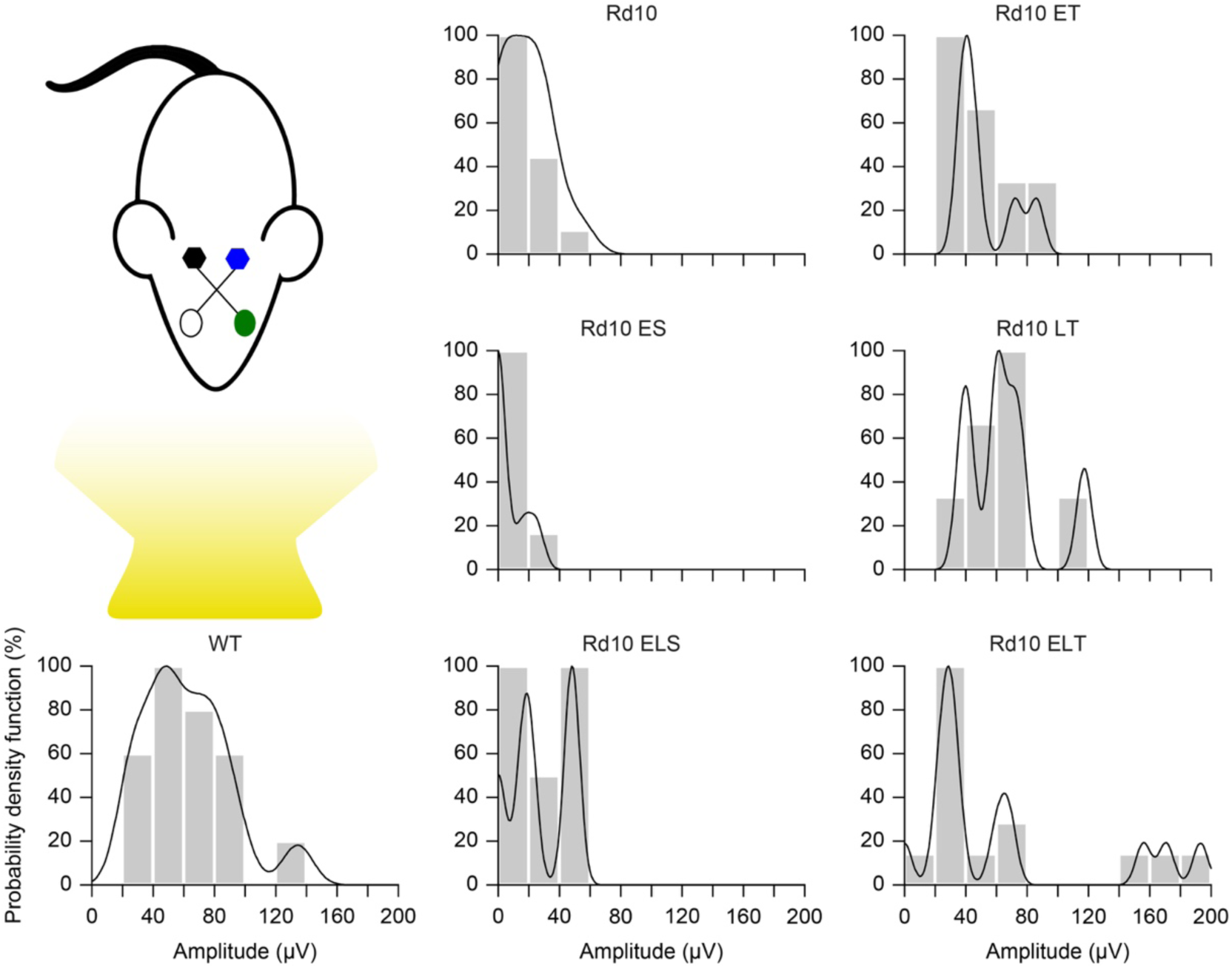
Probability density function. Each panel shows the histogram (20 µV bins) of the VEP peak amplitude and the fitted Kernel distribution for all the experimental groups. For treated (Rd10 ET, Rd10 LT, and Rd10 ELT) and sham treated (Rd10 ES and Rd10 ELS) mice, the data are from recordings in the cortex (black) contralateral to the treated eye (green).

## Discussion

The eye is considered to be a preferential target for the delivery of gene therapies due to its accessibility and immune privilege(Sahel and Roska, 2012). Gene defects affecting photoreceptors cause the vast majority of IRDs; therefore, the development of neuroprotective, gene supplementation, and gene editing therapies has focused primarily on gene transfer to photoreceptors(Lipinski et al., 2015; Smith et al., 2012; Yu et al., 2017). While several retinal cell types can be successfully targeted by different viruses(Colella et al., 2009), the posterior segment of the eye, especially the photoreceptor layer, can be efficiently transduced only by Adeno Associated Viruses (AAVs)(Allocca et al., 2007). Unfortunately, the maximum cargo capacity of these vectors is around 4.7 kb, making them not suitable for the delivery of large genes. CRISPR/Cas9 plasmids are usually bigger than 5 kb and do not fit into AAVs. Hence, to deliver this gene editing tools, it is necessary to either combine more than one AAV vector(Trapani, 2017; Trapani et al., 2014) or use other less safe viral vectors with larger cargo capacity(Rossidis et al., 2018). A smaller Cas9 variant, delivered using AAVs, was recently used to disrupt and thus inactivate the P23H mutated allele in a mouse model of dominant retinitis pigmentosa, but the authors reported a poor cleavage efficiency(Giannelli et al., 2018). Moreover, even though viruses represent the gold standard for gene delivery to the eye, their use does not cease to raise concerns about immunogenicity, long-term safety, and limited possibility for repeated administration(Baum et al., 2006; Thomas et al., 2003; Worgall et al., 2008; Yoshioka et al., 2006). Conversely, non-viral gene delivery strategies(Al-Dosari and Gao, 2009; Niidome and Huang, 2002) permit the multiple administration of large therapeutic agents using less immunogenic and toxic plasmid vectors, but the resulting gene expression is often short-lived(Andrieu-Soler et al., 2007; Bainbridge et al., 2006; Han et al., 2011). This is usually an unappealing characteristic for a clinical application, but it does not represent a concern when editing tools need to be active only for a short period required to correct the sequence of the gene of interest. Among the non-viral delivery strategies, electroporation is one of the most efficient for the introduction of DNA into cells and holds a promising therapeutic potential(Cancedda et al., 2013; Cwetsch et al., 2018; Gehl, 2003; dal Maschio et al., 2012; Szczurkowska et al., 2013; 2016).

In fact, electroporation has been exploited for introducing genetic material and drugs in different tissues and organs and for the treatment of cancer(Cwetsch et al., 2018; Gothelf et al., 2003; Lambricht et al., 2015). According to *in vitro* reports(Hornstein et al., 2016), the size of the plasmid does not have any impact on the transfection efficiency, making the technique suitable for the delivery of large genes that would not fit into AAV viruses. Electroporation has been widely used to study the mouse retina development(Matsuda and Cepko, 2004; 2007; 2008) but is still under investigation for therapeutic purposes. The only reported use of electroporation on the human eye is in the human ciliary muscle(Touchard et al., 2009). Albeit it has not yet been applied to the human retina, it has been shown to successfully target different retinal cell types, both in young(Matsuda and Cepko, 2007; Wang et al., 2014) and adult(Touchard et al., 2012) mice.

In our opinion, electroporation is an attractive alternative method for the delivery of large therapeutic plasmids in the eye. Indeed, CRISPR/Cas9 constructs were also successfully delivered by electroporation in photoreceptor cells to target and disrupt by NHEJ the rhodopsin mutated allele in heterozygous P23H mice(Giannelli et al., 2018; Latella et al., 2016) and in S334ter rats, both model of autosomal dominant retinitis pigmentosa(Bakondi et al., 2016). Nevertheless, loss of function mutations (like the one affecting Rd10 mice) are more difficult to address since the faulty sequence has to be actively edited by HDR to restore the correct gene product and not just disrupted as in the examples cited above.

In this work, retinal electroporation was exploited to deliver a therapeutic DNA mixture to photoreceptor cells. We treated Rd10 mice during photoreceptor development using a CRISPR/Cas9-based gene editing strategy to prevent retinal degeneration and observed preservation of visual functions *in vitro* and *in vivo*, in both subcortical visual-driven behavioural responses (optomotor reflex) and cortical visual responses (VEP) until as late as P90. However, the visual acuity, measured with the optomotor test, eventually declined at the postnatal day 90, even in treated mice. We hypothesise that, since the coverage of the injection is not enough to edit the DNA of all the photoreceptor cells, eventually, also the edited cells succumb to the adverse effect of pro-apoptotic factors released by the non-edited dying cells. Multiple cycles of injections followed by electroporation could solve this issue by allowing the gene editing of a higher number of photoreceptors, especially if performed during the progenitor proliferation period. Notably, we have demonstrated that a repeated treatment (at P3 and P8) is not detrimental for the mice. However, the mouse, especially the pup, is not an ideal model to test this hypothesis, given the tiny size of the eyes: multiple injections and electroporation would damage the eye excessively. Similarly, early intervention on new-born pups (P0-P1) could result in higher editing efficiency.

In conclusion, we provide an example of how CRISPR/Cas9 gene editing can be coupled with electroporation for therapeutic purpose, along with a discussion of the limitations that need to be overcome to translate this approach to the clinics. Issues related to safety of retinal electroporation in large animals, repeatability of the treatment, transfection efficiency, retinal coverage, gene expression levels, and disease stage at the age of injection still need a careful investigation to improve the therapeutic benefits of CRISPR/Cas9-based gene editing strategies. On the other hand, the ease of design of the CRISPR/Cas9 gene editing systems makes them easily tailorable for several mutations in perspective of a patient-specific therapy. This concept applies particularly when there are small differences in the DNA sequence, as in the case of autosomal recessive mutations. Our non-viral delivery approach has two main advantages compared to previous reports in small animal models in which retinal degeneration was prevented by viral-mediated delivery of Cas9. First, in perspective of a possible clinical application, it circumvents possible safety issues deriving from viral-based gene therapy. Secondly, plasmid vector delivery by electroporation will result in transient expression of Cas9, therefore limiting the possible occurrence of off-target activity of the nuclease after long-term expression.

## Methods

### Construct Design and Cloning

The online CRISPR Design Tool (http://tools.genome-engineering.org) was used to design the gRNAs targeting the mouse gene *Pde6b* at the level of the mutation C1678T. The sequence of the gene was used as input, and the first three best scoring gRNAs were selected. gRNA #1 and gRNA #3 flanked the mutation C1678T (mapping respectively upstream and downstream the mutation), while gRNA #2 and its Rd10-mutated counterpart gRNA #4 mapped on the mutation. The gRNA sequences (#n) are the following: gRNA #1, gtggtaggtgattcttcgat; gRNA #2, tgaagccgtggc**g**ccagttg; gRNA #3, tctgggctacattgaagccg; gRNA #4, tgaagccgtggc**a**ccagttg. The gRNA #2 and #4 differ only for one base (in bold). The oligonucleotides to generate the gRNAs (Integrated DNA Technologies) were annealed *in vitro* and cloned in the BbsI sites of the pSpCas9(BB)-2A-GFP plasmid (#48138, Addgene). The original CBh promoter in pSpCas9(BB)-2A-GFP plasmid was then replaced with the CAGGs promoter from pCAGGs–mCherry (#41583, Addgene). The obtained pCAGGs-Cas9(BB)-2A-GFP-gRNA plasmid was subsequently used for the *in vitro* experiments. To design the single-stranded oligodeoxynucleotide (ssODN) to use as repair templates, we took advantage of a BanI restriction site in the target sequence to develop a screening assay that allowed us to distinguish between the edited and the non-edited sequences. The BanI restriction site (GGYRCC, where Y = C or T and R = A or G) maps just downstream to the C1678T mutation and is present both in the WT (GGCGCC) and the Rd10 (GGTGCC) *Pde6b* sequence. We designed individual ssODN repair templates for each gRNA (**Tab. 1**) in order to introduce a silent mutation in the corresponding gRNA PAM sequence (NGG) to avoid Cas9 mediated re-processing of the edited DNA strand. Moreover, the repair ssODNs (Integrated DNA Technologies, Ultramer™ DNA oligo) were designed to restore the codon coding for the Arginine (mutated into a Cystein in the Rd10 mice, R560C) and concomitantly destroy the BanI restriction site (TGC to AGA). Since the gRNA #2 and #4 differ only by one base, they share the same ssODN.

**Table 1.**
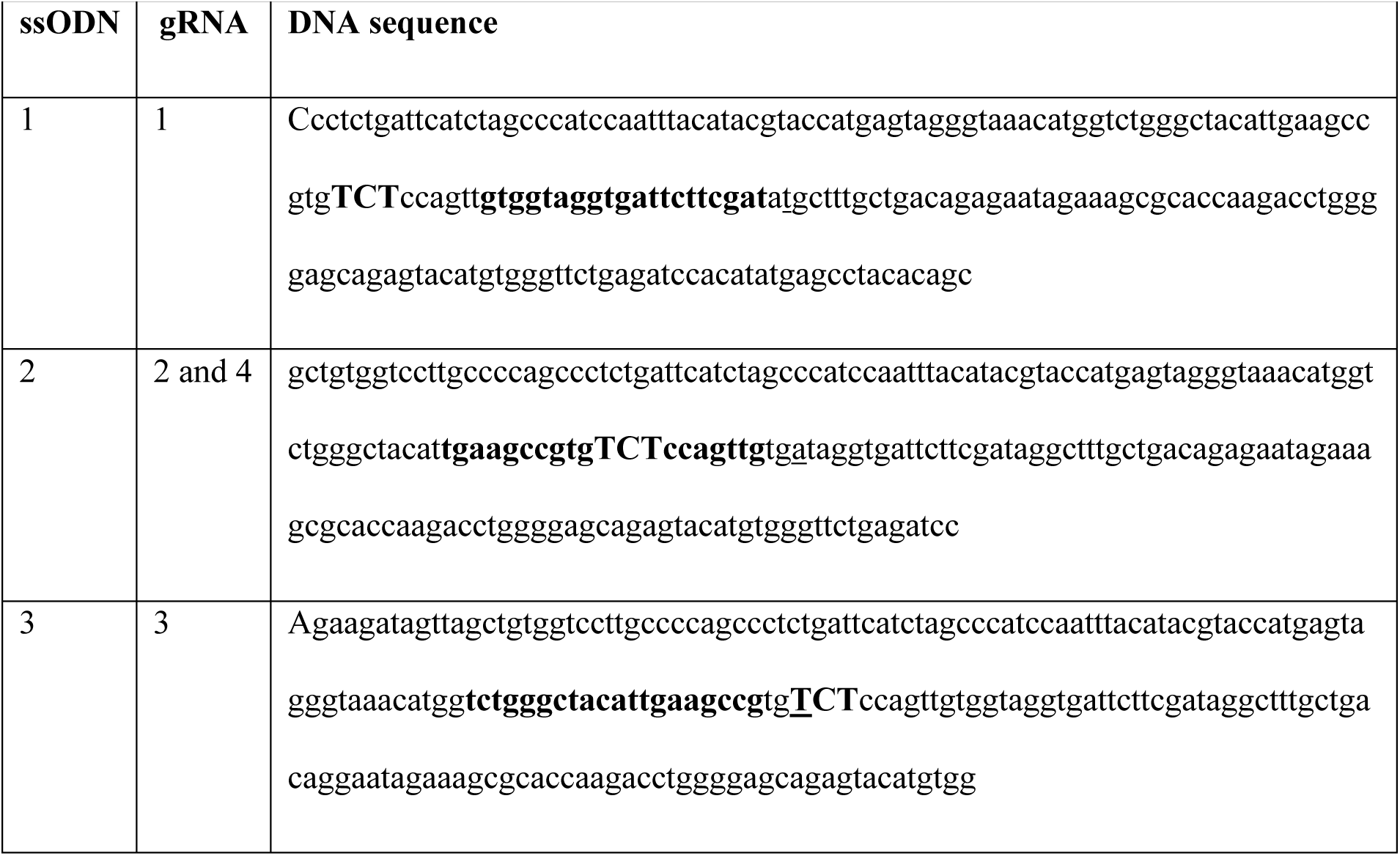
ssODNs coupled with each gRNA. The sequences in bold represent the gRNAs and the ones in bold capital letters represent the edited bases that restore the Arginine and delete the BanI restriction site. The bases underlined are those mutated in the PAM sequence. The ssODNs are antisense to the Pde6b sequence.

### N2A Cell Culture and Transfection

Mouse N2A cells (ATCC® CCL-131(tm)) were cultured in Dulbecco’s minimum essential medium (DMEM, Life Technologies) supplemented with 10 % foetal calf serum (Life Technologies), 1 % L-glutamine, 100 U ml^-1^ penicillin, and 100 mg ml^-1^ streptomycin (Biowhittaker-Lonza). Cells were maintained at 37 °C in a 5 % CO2 humidified atmosphere. The cells were transfected with Fugene 6 (Roche). The day before transfection 5 x 10^5^ N2A cells were plated on 6 cm plates. The medium was replaced with fresh medium 1 h before the transfection. The DNA/Fugene mix (ratio 1:2) was prepared in Optimem medium (Life Technologies). N2A cells were co-transfected with 1.5 μg of pCAGGs-Cas9(BB)-2A-GFP-gRNA #(n) and 2.2 μg of the associated repair template. Cells plated on different wells were transfected with different gRNAs. Cells were incubated at 37 °C in a 5 % CO_2_ humidified atmosphere for 48 h following transfection, then detached using Trypsin-EDTA 0.25 % (Sigma-Aldrich), and prepared for FAC-sorting.

### Preparation of Neurospheres and Nucleofection

Primary cultures of NSC were prepared from WT and Rd10 mice(Pacey et al., 2006). P2 mice were decapitated, and the brain was removed from the skull. The cortex and the hippocampus were isolated and cut in small cubes in the tissue dissection solution (in mM): 124 NaCl, 5 KCl, 3.2 MgCl_2_, 26 NaHCO_3_, 10 glucose, and 0.1 CaCl_2_ (Sigma-Aldrich). An enzyme mix was dissolved in 30 ml of tissue dissection solution and added: trypsin 0.04 g, Type 1-S Hyaluronidase 0.02 g, and kynurenic acid 0.004 g (Sigma-Aldrich). The tissue was incubated for 40 min in a water bath at 37 °C and triturated with a Pasteur pipette every 20 min. After centrifugation, the enzyme mix was removed and the trypsin inhibitor (Sigma-Aldrich), dissolved in serum-free medium (SFM) at the concentration of 1 mg ml^-1^, was added. The tissue was then triturated and incubated in the water bath for an additional 10 min. After centrifugation, the tissue was resuspended in SFM containing: DMEM/F12 (Life Technologies), 20 ng ml^-1^ EGF (Peprotech), 20 ng ml^-1^ FGF (Peprotech), 2 % v/v B-27 (Life Technologies), 1.83 µg ml^-1^ Heparin (Sigma-Aldrich), 1 mM Putrescine (Sigma-Aldrich), 2 µM Progesterone (Sigma-Aldrich), 10 µg ml^-1^ ITSS (Sigma-Aldrich), 6 mg ml^-1^ glucose (Sigma-Aldrich), and 1 % Pen/Strep (Life Technologies). Then, the tissue was triturated to obtain a single-cell solution. The cells were counted with the vital dye Trypan blue (Sigma-Aldrich) and then plated at 100.000 cells in each well of a 12-well non-coated plate. We obtained neurospheres that were maintained in SFM at 37 °C in a 5 % CO2 humidified atmosphere and passed 1:3 for three times a week. After 3 to 4 passages cell were electroporated *via* Nucleofection with the AMAXA nucleofection device (LONZA). The Neurospheres were dissociated with Accutase (Sigma-Aldrich) and 5 x 10^6^ NSCs were electroporated with 2 μg of pCAGGs-Cas9(BB)-2A-GFP-gRNA and 2 μl of repair template (10 μM) following the protocol suggested by the manufacturer. Cells were then incubated at 37 °C in a 5 % CO_2_ humidified atmosphere for 30 hours, dissociated with Accutase, and GFP-positive cells isolated by FACS.

### Restriction analysis

Cells in Hibernate-A medium were filtered (Life Technologies) and FACS-isolated with a FACSAria (BD-Biosciences). GFP positive cells were collected in a tube containing PBS + FBS 2 %. The gDNA of the sorted cells (both N2A and NSCs) was extracted with the Genomic DNA(tm) - Tissue MiniPrep kit (Zymo Research) following the protocol of the manufacturer for cell suspensions. The DNA was eluted in 30 µl of DNAse-free water and concentration measured at 260 nm with an ND1000 Nanodrop spectrophotometer (Thermo Scientific). 125 ng of purified gDNA was used for PCR amplification. The following primers (Sigma-Aldrich), mapping outside the ssODN sequence were used to amplify a region of ≈ 700 bps containing the edited region of the *Pde6b* gene: tttctgctcacaggccacat (forward) and gctccagaaggcagtggtta (reverse). The DNA fragment obtained by amplification was purified with the PCR purification kit (QIAGEN) and quantified as above. For the restriction analysis of PCR products, 300 ng of DNA was digested with 5 units of BanI enzyme for 1 hour in 25 µl total reaction volume. The digestion of the PCR fragment obtained amplifying unedited gDNA with the BanI restriction enzyme generated two fragments of 470 and 230 bps respectively that were resolved on 2 % agarose gel. The PCR fragments obtained amplifying the edited gDNA could not be digested by the BanI enzyme, thus leaving the undigested 700 bps fragment on an agarose gel. The optical density of the 700-bps band was measured using the gel tool of ImageJ.

### Plasmids and DNA preparation for in vivo delivery

The nanoplasmids expressing eGFP and Cas9/GFP were purchased from Nature Technology, and the template repair was purchased from Integrated DNA Technologies. The gRNA was cloned into the pSPgRNA plasmid (#47108, Addgene). All the components used for the *in vivo* experiments are specified in **Table 2**.

**Table 2.** Full name, size, and origin of all the components used in the in vivo experiments. The concentrations were adjusted in order to have the same number of copies of guide-coding and Cas9-coding plasmids, taking into account the relative number of base pairs. The repair ssODN concentration is similar to what previously described for CRISPR-Cas9 editing systems.

| Component | Name | Length | Supplier |
| --- | --- | --- | --- |
| eGFP plasmid | NTC9385R-eGFP | 2391 bp | Nature Technology |
| Cas9 plasmid | NTC9385R-CAGCas9-T2A-GFP | 6500 bp | Nature Technology |
| gRNA plasmid | pSPgRNA | 3000 bp | Addgene |
| Repair template | ssODN | 200 nucleotides | IDT |

The following are the specific solutions used for each experiment (all of them were prepared in PBS with the addition of 0.1 % Fast green for the visualisation of the injection):

-eGFP preparation: eGFP-coding plasmid (1 µg µl^-1^).
-Cas9 preparation: Cas9-coding plasmid (1 µg µl^-1^) + gRNA-coding plasmid (0.45 µg µl^-1^) + repair ssODN (2 µl µg of Cas9^-1^).
-Sham preparation: Cas9-coding plasmid (1 µg µl^-1^) + repair ssODN (2 µl µg of Cas9^-1^).
-Cas9 + eGFP (1.5:1) preparation: Cas9-coding plasmid (1.5 µg µl^-1^) + gRNA-coding plasmid (0.8 µg µl^-1^) + eGFP-coding plasmid (0.9 µg µl^-1^) + repair ssODN (2 µl µg of Cas9^-1^).

### Animal handling

Animal experiments were performed according to the animal authorizations VD3044 issued by the Service de la consommation et des Affaires vétérinaires (SCAV) of the Canton de Vaud (Switzerland), GE3217 issued by the Département de l’emploi, des affaires sociales et de la santé (DEAS), Direction générale de la santé of the Republique et Canton de Genève (Switzerland), and 726/2015-PR issued by the Italian Ministry of Health. Male and female mice pups at P3 and P8 and adult mice at 1, 2, and 3 months of age from a homozygous colony of B6.CXB1-*Pde6b*^*rd10*^/J mice (The Jackson Laboratory) were used for the experiments. C57BL/6J mice (Charles River) were used as control group. All animals were kept in a 12 h day/night cycle with access to food and water ad libitum. All pups were kept with the mother until weaning, except for the time necessary to perform the subretinal injection. All the experiments were carried out during the day cycle.

### Subretinal injection and electroporation

Subretinal injections were performed in mice pups at P3, P8, or both. The pups were anaesthetised using isoflurane (0.8-1.5 l min^-1^ at 3 %) in an induction box, then placed onto a sterile paper towel under a microscope; the anaesthesia was maintained with isoflurane (0.8-1.5 l min^-1^ at 2 %), and the temperature was maintained at 37 °C with a heating pad. The skin over the eyelid was disinfected with Betadine, and a sterile 30-G needle was used to cut the skin on the mark of the future eyelid aperture. The skin was gently pushed to the side with a pair of sterile forceps to expose the eyeball. A glass capillary (ORIGIO) backfilled with the DNA solution was insert into the subretinal space, maintaining a 45° inclination to the surface of the eye. The DNA solution was then injected into the subretinal space for 3 seconds at 300 hPa using an automatic injector (Eppendorf). Two injections were performed in the following directions: dorsal to nasal and ventral to nasal. Immediately after the DNA injection, an electric field was applied to the area using a P5 tweezer electrode (Sonidel) pre-soaked in PBS. The positive terminal was attached to the sclera of the injected eye, while the other side of the tweezer (negative terminal) was placed on the not-injected eye. The pulses were delivered using a CUY21SC electroporator (Sonidel). A conductive gel was placed between the electrode plate and the eye to maximise the conductivity and minimise burns on the cornea. Two square pulses of 5 ms at 100 V were applied with 0.1 Hz frequency (poring pulses), followed by five pulses of 50 ms at 30 V with 1 Hz frequency (transfer pulses). After the procedure, the eyelid was closed gently with a cotton swab, and the pup was placed onto a heating pad at 37 °C until fully recovered, then returned to the mother. In all the groups the injection was performed unilaterally in order to keep the other eye as an internal control. The pups were treated daily during the first-week post-surgery with Tobradex Eye Drops (tobramycin 0.3 % and dexamethasone 0.1 %) on the operated eye.

### Retina sections and wholemounts

Mice for retina sections and wholemounts were electroporated at P3 or P8 with the eGFP construct. After euthanasia by CO_2_ inhalation, the eyes of the mice were extracted from the ocular cavity using forceps, washed in PBS, and fixed in 4 % PFA overnight. For wholemount preparation, the retina was extracted and cut in 4 points in order to flat it on a microscope slide. For section preparation, the samples were cryoprotected in sucrose 30 % and frozen in optimal cutting temperature compound (Tissue-Tek®). 20 µm thick sections of the retina were obtained using a Histocom cryostat (Thermo Scientific) and placed on microscope slides. The sections and the wholemounts were washed in PBS, permeabilised with PBS + Triton 0.1 % (Sigma-Aldrich), counterstained with DAPI 1:300 (Sigma-Aldrich), and mounted for imaging with Fluoromount (Sigma-Aldrich).

### Droplet-digital PCR

The eyes of P6 mice electroporated at P3 were enucleated, and the retina was immediately isolated in ice-cold PBS and quickly inspected under a fluorescence microscope to verify eGFP expression. The gDNA was extracted using the Genomic DNA™ -Tissue MiniPrep kit (Zymo Research) following the protocol of the manufacturer for solid tissues. The DNA was eluted in 30 µl of DNase-free water. To avoid possible false-positive signals in ddPCR from unintegrated ssODN repair template, we optimised a nested-ddPCR assay. We first pre-amplified from extracted gDNA by conventional PCR a fragment of ≈700 bps containing the edited region of the *Pde6b* gene with primers mapping outside the ssODN sequence (same as for the BanI restriction assay). The amplified DNA fragment was purified and quantified as above. Next 2.5 fg of the purified template (corresponding to ≈3500 copies of target DNA) was used in the ddPCR assay with internal primers (Fwd: CAGCAAAGCCTATCGAAGAATCA; Rev: CATGGTCTGGGCTACATTGAAG) and detected with an edited-specific TaqMan® probe (FAM-TATCACAACTGGAGACAC-MGB) and an unedited-specific TaqMan® probe (VIC-TACCACAACTGGTGCCA-MGB). Reactions were assembled with ddPCR(tm) Supermix for Probes (Bio-Rad Laboratories) and partitioned into nanoliter-sized droplets with QX200 Droplet Generator (Bio-Rad Laboratories). After PCR thermal cycling, droplets for each sample were individually read on a QX200 Droplet Reader (Bio-Rad Laboratories) and assigned as positive or negative based on fluorescence amplitude.

### Recordings of microelectroretinograms

P60 mice electroporated at P3 were dark-adapted overnight before tissue collection. All procedures were performed under dim red light. Retinas were explanted after euthanasia by intraperitoneal injection of sodium pentobarbital (150 mg kg^-1^). The retinas were dissected in carboxygenated (95 % O_2_ and 5 % CO_2_) Ames’ medium (A1420, Sigma-Aldrich). After dissection of the sclera, the retina was detached from the pigment epithelium, and the vitreous humour was removed. The retina was then cut into pieces (approximately 5 mm^2^), attached to a filter paper, and transferred onto the MEA (256MEA200/30iR-ITO; Multi Channel Systems) with the ganglion cell layer facing the electrodes. Explanted retinas were continuously superfused with carboxygenated Ames’s medium at 32 °C. Data acquisition, amplification, and digitalisation were performed with a recording system (USB-MEA256-System; Multi Channel Systems) placed on the stage of an inverted Ti-E microscope (Nikon Instruments). The microscope was equipped with a dichroic filter (FF875-Di01-25×36; Semrock) and a 4x objective (diameter of the illumination spot 5.5 mm; CFI Plan Apochromat Lambda). Light stimuli were provided by an attached Spectra X system (Emission filter 560/32; Lumencor). Ten consecutive pulses of 4 ms and 0.5 mW mm^-2^ were delivered at a repetition rate of 1 Hz. The extracellularly recorded signals were digitalised and stored for offline analysis. Data filtering and spike sorting were performed using the MC_Rack software (Multi Channel Systems). The presence of spontaneous spiking activity was assessed (filter 300-3000 Hz, sampling rate of 25 kHz) to ensure the viability of the retinal explant. Only retinas showing spontaneous activity in at least one channel when placed on the MEAs were selected for recordings, and each responding channel was treated as an independent unit. To detect µERGs, the signal was filtered from 0.5 to 100 Hz and digitalised at 10 kHz. The prominence of the µERG a-wave was computed for each channel in MATLAB (MathWorks).

### Measurement of the visual acuity

The Optomotor system (Cerebral Mechanics) was used for the measurement of the visual acuity. Control and treated mice were habituated for 5 min placing them in the centre of the virtual arena the day before the beginning of the test. The day of the test, each mouse was placed on the platform, and the program started. The mouse in the arena was presented with a grating stimulus rotating in either direction, and the operator had to decide if the mouse was tracking or not the rotating stimulus with a movement of the head in the same direction of the rotation. The program uses a built-in algorithm based on a staircase method to evaluate the visual threshold of the two eyes independently. The performance of any single mouse was assessed during three subsequent days, and the resulting average was considered the value of the mouse visual threshold. Mice were tested at P30, P60, and P90.

### Electrode implantation and recording of visually evoked potentials

Before the surgical procedures and the recording sessions, the mice were anaesthetised with isoflurane (0.8-1.5 l min^-1^ at 4 % for induction and 0.8-1.5 l min^-1^ at 1.5 % for maintenance). Analgesia was administered by subcutaneous injection of Buprenorphine (Temgesic, 0.1 mg kg^-1^) followed by subcutaneous injection of a mix composed by lidocaine (6 mg kg^-1^) and bupivacaine (2.5 mg kg^-1^) with a 1:1 ratio. The depth of anaesthesia was assessed with the pedal reflex, and artificial tears were used to prevent the eyes from drying. The temperature was maintained at 37 °C with a heating pad during both surgical and recording sessions. For electrode implantation, anaesthetised P60 mice mounted on a stereotaxic apparatus. The skull was exposed for the visualisation of lambda, and the skin was pulled on the side. Two screw electrodes were implanted 3 mm lateral to lambda over the left and right visual cortices. A reference electrode was placed in the rostral side of the cranium, outside of the visual cortex. The electrodes were fixed using dental cement. The screws were then left in place for 30 more days. The surgery was performed in advance in order to let the electrodes to stabilise. For recordings, all the procedures were performed under dim red light. P90 mice were dark-adapted overnight, anaesthetised, and mounted on a stereotaxic frame. The pupils were dilated with a drop of Atropine 1 %, and a needle electrode was placed subcutaneously in the dorsal area near the tail as ground. The recordings were acquired simultaneously in two channels connected to the two electrodes implanted on both visual cortices. Three light flashes (4 ms, 10 cd s m^-2^, interleaved by 2 min) were delivered with a Ganzfeld stimulator (BM6007IL, Biomedica Mangoni) positioned close to the mice and the corresponding visually evoked cortical potentials were amplified, filtered (0.1 – 500 Hz), and digitalized for 1000 ms (50 ms pre-stimulus and 950 ms post-stimulus) at 2 kHz (BM623, Biomedica Mangoni). The data were analysed using MATLAB (MathWorks).

### Statistical analysis and graphical representation

Statistical analysis and graphical representation were performed with Prism (GraphPad Software Inc.). The normality test (D’Agostino & Pearson omnibus normality test) was performed in each dataset to justify the use of a parametric (t-test and One-Way ANOVA) or non-parametric (Kruskal-Wallis and Mann-Whitney) test. The fitting of the VEPs was performed with the non-parametric Kernel distribution in MATLAB. In each figure p-values were represented as: * p < 0.05, ** p < 0.01, *** p < 0.001, and **** p < 0.0001.

### Data availability

The authors declare that all other relevant data supporting the findings of the study are available in this article and its Supplementary Information file. Access to our raw data can be obtained from the corresponding author upon reasonable request.

## Acknowledgements

We thank Elodie Meyer at the Medtronic Chair in Neuroengineering for her help with retinal histology, Aurelién Lacombe at the Wyss Center for Bio and Neuroengineering for his support on molecular biology, Ennio Albanesi at the Istituto Italiano di Tecnologia for FACS isolation of cells, and Dr. Sui Wang and Prof. Constance Cepko at Harvard Medical School for their help with subretinal electroporation. This work was supported by École Polytechnique Fédérale de Lausanne, Medtronic, 2014 Global Ophthalmology Awards Program (GOAP) by Bayer HealthCare to D.G., Compagnia di San Paolo (Italy) grant 9734 to A.C., and the European Research Council (ERC) under the European Union’s Horizon 2020 research and innovation program (grant agreement No 725563) to L.C.

## Author Contributions

P.V. performed all the experiments and wrote the manuscript. L.E.P. designed the gene editing strategy, generated the plasmids, and performed in vitro validation of the plasmids. N.A.L.C. performed experiments with explanted retinas. T.M. contributed to in vivo electrophysiology. M.P. contributed to the setting of pilot behavioural experiments. A.C. designed the study, designed the gene editing and screening strategies, performed the ddPCR and wrote the manuscript. L.C. designed the study. D.G. designed the study, led the whole project, and wrote the manuscript. All the authors read, edited, and accepted the manuscript.

